# Leveraging longitudinal diffusion MRI data to quantify differences in white matter microstructural decline in normal and abnormal aging

**DOI:** 10.1101/2023.05.17.541182

**Authors:** Derek B. Archer, Kurt Schilling, Niranjana Shashikumar, Varuna Jasodanand, Elizabeth E. Moore, Kimberly R. Pechman, Murat Bilgel, Lori L. Beason-Held, Yang An, Andrea Shafer, Luigi Ferrucci, Shannon L. Risacher, Katherine A. Gifford, Bennett A. Landman, Angela L. Jefferson, Andrew J. Saykin, Susan M. Resnick, Timothy J. Hohman, Alzheimer’s Disease Neuroimaging Initiative

**Affiliations:** Vanderbilt Memory and Alzheimer’s Center, Vanderbilt University School of Medicine, Nashville, TN, USA; Vanderbilt Genetics Institute, Vanderbilt University Medical Center, Nashville, TN, USA; Laboratory of Behavioral Neuroscience, National Institute on Aging, National Institutes of Health, Baltimore, MD, USA; Indiana University School of Medicine, Indianapolis, IN, USA; Indiana Alzheimer’s Disease Research Center, Indianapolis, IN, USA; Vanderbilt University Institute of Imaging Science, Vanderbilt University Medical Center, Nashville, TN, USA; Department of Biomedical Engineering, Vanderbilt University, Nashville, TN, USA; Department of Electrical and Computer Engineering, Vanderbilt University, Nashville, TN, USA; Department of Medicine, Vanderbilt University Medical Center, Nashville, TN, USA; Department of Radiology & Radiological Sciences, Vanderbilt University Medical Center, Nashville, TN, USA; National Institute on Aging, Baltimore, MD, USA

**Author notes:** **Consent Statement:** All participants provided informed consent in their respective cohort studies. **Corresponding Author:** Derek B. Archer, PhD Department of Neurology Vanderbilt University Medical Center 1207 17^th^ Ave South, Suite 204 Nashville, TN 37212, USA.

**Keywords:** White matter, diffusion MRI, harmonization, free-water, aging

## Abstract

**INTRODUCTION:** It is unclear how rates of white matter microstructural decline differ between normal aging and abnormal aging.

**METHODS:** Diffusion MRI data from several well-established longitudinal cohorts of aging [Alzheimer’s Neuroimaging Initiative (ADNI), Baltimore Longitudinal Study of Aging (BLSA), Vanderbilt Memory & Aging Project (VMAP)] was free-water corrected and harmonized. This dataset included 1,723 participants (age at baseline: 72.8±8.87 years, 49.5% male) and 4,605 imaging sessions (follow-up time: 2.97±2.09 years, follow-up range: 1–13 years, mean number of visits: 4.42±1.98). Differences in white matter microstructural decline in normal and abnormal agers was assessed.

**RESULTS:** While we found global decline in white matter in normal/abnormal aging, we found that several white matter tracts (e.g., cingulum bundle) were vulnerable to abnormal aging.

**CONCLUSIONS:** There is a prevalent role of white matter microstructural decline in aging, and future large-scale studies in this area may further refine our understanding of the underlying neurodegenerative processes.

**HIGHLIGHTS:**

- Longitudinal data was free-water corrected and harmonized
- Global effects of white matter decline were seen in normal and abnormal aging
- The free-water metric was most vulnerable to abnormal aging
- Cingulum free-water was the most vulnerable to abnormal aging

## Introduction

While white matter microstructural decline has been well-characterized in normal aging^1–15^, fewer studies have focused on how these aging patterns differ in participants with neurodegenerative disease. Understanding how aging patterns differ in normal and abnormal aging would further our understanding into which white matter tract vulnerabilities exist in disease. In Alzheimer’s disease (AD) – the most prevalent neurodegenerative disease in abnormal aging^16^ -- it is unclear which white matter tracts decline in the aging process. Given that white matter has been strongly implicated to play a role in the development of AD^17, 18^, paired with recent evidence that *apolipoprotein E-*ε4 status leads to cholesterol deposits in oligodendrocytes resulting in demyelination^19^, it is pivotal that large-scale studies determine how white matter microstructural decline differs between normal aging and aging in association with cognitive impairment, including AD. Moreover, it is crucial that we understand which white matter tracts are most vulnerable to AD.

In normal aging, initial neuroimaging investigations studying white matter microstructural decline primarily used region of interest (ROI) approaches, with particular emphasis on the genu and splenium of the corpus callosum^7, 20^. One study used T2-weighted MR images to calculate the transverse relaxation rate – a measure sensitive to tissue microstructure – within the genu and splenium in 252 participants ranging in age from 19-82 years. They found that the genu demonstrated a strong inverted U-function, whereby subtle increases in the relaxation rate were exhibited early in life with a drastic decline beginning in the third decade of life. In the splenium, there was a less robust linear association with age, such that higher age was associated with lower relaxation rates^20^. With respect to diffusion MRI, a seminal study evaluated whether aging was associated with abnormal fractional anisotropy (FA) within several ROIs. They found that the most significant age-related changes in FA were within the genu, posterior limb of the internal capsule, and the frontal white matter^7^. These results, in conjunction with the release of easy-to-use ROI and tractography atlases^21, 22^, invigorated further research into how normal aging is associated with white matter microstructural decline. The prevailing hypothesis from these studies is that late-myelinating tracts, such as those projecting from the frontal white matter, are particularly vulnerable in normal aging^3, 15^.

Prior literature has strongly implicated that participants with AD exhibit global white matter differences, with pronounced microstructural decline in the temporal lobe^4, 14, 23–29^. Recent evidence, however, has drastically enhanced our knowledge in this space^30, 31^. One cross-sectional study (n=405) quantified cholinergic pathway and cingulum bundle microstructure along the AD continuum, and found these tracts were vulnerable even at the earliest stages of disease (i.e., subjective cognitive decline)^30^. A recently published longitudinal study of aging (n=250) compared rates of white matter microstructural decline between cognitively unimpaired individuals with individuals with subsequent memory impairment^31^. They found that individuals with subsequent memory impairment had more rapid rates of white matter decline in the splenium of the corpus callosum and inferior frontal occipital fasciculus. Together, these studies suggest that there is widespread white matter vulnerability associated with cognitive impairment and AD, with pronounced effects in the limbic, association, and occipital transcallosal tracts.

While the previously mentioned studies in normal aging and AD have been foundational to our understanding of white matter microstructural decline with age, the status quo in a large majority of research has been to use long-standing white matter tractography atlases. Recently developed tractography templates, however, have enhanced coverage and improve spatial specificity which may further our understanding of the aging brain^32–35^. For example, we have created tractography templates of the sensorimotor tracts, transcallosal projections, and medial temporal lobe projections which have already demonstrated utility in a variety of neurodegenerative disorders^17, 34, 35^. Incorporating these spatially precise white matter tractography templates will give us unparalleled insight into the patterns of white matter microstructural decline that differ in normal aging and AD.

In addition to new tractography templates, advanced post-processing techniques now allow for researchers to correct for well-known confounds, such as partial volume, in diffusion MRI acquisitions^36, 37^. For example, neurite orientation dispersion and density imaging (i.e., NODDI) can be conducted on multi-shell diffusion MR images to estimate the orientation dispersion index, intracellular volume fraction, and the isotropic volume fraction^37^. While this advanced technique is the preferred method to evaluate multiple compartments in diffusion MR images, many long-standing cohorts of aging have predominantly single-shell diffusion MR images available; thus, retrospective analyses using these cohorts require alternative post-processing techniques. Fortunately, there are established techniques which allow researchers to correct for partial volume effects in single-shell diffusion MR images. One technique, developed by *Pasternak et al.,* uses a bi-tensor model to characterize an extracellular component [i.e., the free-water (FW) component], which reflects the amount of unrestricted water movement within a voxel^36^. This extracellular component is then removed from the image and FW-corrected intracellular metrics can be quantified, including FA (FA_FWcorr_), axial diffusivity (AxD_FWcorr_), radial diffusivity (RD_FWcorr_), and mean diffusivity (MD_FWcorr_). This method has already been used to study several neurodegenerative disorders, including chronic stroke^38^, essential tremor^39^, Parkinsonism^34, 40–42^, schizophrenia^43^, and Alzheimer’s disease^17, 44^. With respect to normal aging, a recent study used FW correction to evaluate a cohort of 212 participants ranging from 39-92 years^45^. In this study, they found that the effect of normal aging was mitigated in intracellular metrics after FW correction, suggesting that these metrics are biased by the extracellular component and could be due to larger interstitial spaces and/or inflammation. They further concluded that the aging effect on FW was consistent with the anterior-posterior hypothesis, whereby anterior regions of the brain had higher FW compared to posterior regions. There has yet to be a large-scale analysis leveraging FW correction to understand the differences in white matter microstructural decline between normal and abnormal aging.

In the present study, we used single-shell diffusion MRI data from the Alzheimer’s Disease Neuroimaging Initiative (ADNI), Baltimore Longitudinal Study of Aging (BLSA), and Vanderbilt Memory & Aging Project (VMAP) cohorts. In total, this study used data from 1,723 participants across 4,605 imaging sessions. All diffusion MRI data were preprocessed with an identical pipeline and conventional and FW-corrected microstructural values were quantified within 48 white matter tracts spanning the association (n=3), limbic (n=7), projection (n=9), and transcallosal (n=29) tracts. Given that there were different scanners, sites, and protocols, we harmonized all microstructural values using the *Longitudinal ComBat* technique and these harmonized values were used to study the effect of aging on white matter. This harmonized data was then used to: (1) determine how normal and abnormal aging are associated with white matter microstructural decline, (2) quantify the interaction between normal and abnormal aging on white matter microstructural decline, and (3) investigate which white matter tracts are most vulnerable to abnormal aging. We hypothesized that limbic, prefrontal transcallosal, and occipital transcallosal tracts would exhibit the greatest differential rates of white matter microstructural decline between normal and abnormal aging.

## Methods

### Study Cohort

The present study used data from 3 well-established cohorts of aging. The largest cohort was the Neuroimaging substudy of the BLSA^46^ – behavioral assessment in this cohort began in 1994 and included dementia-free participants aged 55-85 years. From 2006-2018, BLSA MRI data was collected on a 1.5T scanner, and in 2009 this cohort was expanded to include participants aged 20 and older and 3T MRI data collection began on a single scanner. Data from the BLSA cohort are available upon request by a proposal submission through the BLSA website (www.blsa.nih.gov). Another cohort leveraged in this study was the well-known ADNI (adni.loni.usc.edu) cohort^47^ – this cohort was launched in 2003 as a public-private partnership, led by Principal Investigator Michael W. Weiner, MD. The primary goal of ADNI has been to test whether serial magnetic resonance imaging (MRI), positron emission tomography (PET), other biological markers, and clinical and neuropsychological assessment can be combined to measure the progression of mild cognitive impairment (MCI) and early AD. The final cohort used in this study was VMAP^48^ – data collection for VMAP began in 2012 and includes participants aged 60+ years who are considered cognitively unimpaired or have mild cognitive impairment. Data from the VMAP cohort can be accessed freely following data use approval (www.vmacdata.org). Within each cohort, informed consent was provided by all participants and all studies were conducted in accord with the Declaration of Helsinki. For each cohort, several demographic and clinical covariates were required for inclusion, including age, sex, educational attainment, race/ethnicity, apolipoprotein E (*APOE*) haplotype status (ε2, ε3, ε4), and cognitive diagnosis [cognitively unimpaired (CU), mild cognitive impairment (MCI), Alzheimer’s disease (AD)]. Participants aged 50+ years at baseline were included in the present study. Cognitive diagnosis across each participant’s imaging sessions were evaluated to determine if they were “normal” or “abnormal” agers. In total, this study included 1,723 participants across 4,605 imaging sessions. Participants were considered normal agers if they had a CU diagnosis across all imaging sessions, whereas participants with any non-CU diagnosis across any imaging session were considered abnormal agers. Sample sizes, demographic information, and health characteristics for each cohort can be found in **Table 1**, and parameters for each MRI acquisition included in this study can be found in **Supplemental Table 1**.

**Table 1.** – Demographic and Health Characteristics

| Measure | Cohort |  |  | p-value |
| --- | --- | --- | --- | --- |
|  | ADNI | BLSA | VMAP |  |
| Cohort Characteristics |  |  |  |  |
| Number of participants | 706 | 721 | 296 | - |
| Total number of sessions | 1,859 | 1,858 | 888 | - |
| Average number of visits | 4.89 (2.41) | 4.27 (1.84) | 3.78 (0.54) |  |
| Longitudinal follow-Up (yrs)* | 1.97 (1.85) | 3.94 (2.13) | 3.01 (1.47) | <0.001 |
| Demographics and Health Characteristics |  |  |  |  |
| Age at baseline (yrs) | 74.03 (7.72) | 71.33 (10.21) | 73.32 (7.26) | <0.001 |
| Sex (% male) | 51.42 | 44.24 | 57.77 | <0.001 |
| Education (yrs) | 16.29 (2.62) | 17.02 (2.38) | 15.81 (2.68) | <0.001 |
| Race (% Non-hispanic white) | 91.64 | 65.74 | 92.91 | <0.001 |
| APOE-ε4 (% positive) | 42.21 | 26.91 | 35.81 | <0.001 |
| APOE-ε2 (% positive) | 8.92 | 17.06 | 15.20 | <0.001 |
| Cognitive status at baseline (% cognitively unimpaired) | 46.74 | 98.34 | 56.76 | <0.001 |
| Longitudinal cognitive status (normal/abnormal) | 395/311 | 676/45 | 168/128 | <0.001 |

Values denoted as mean (standard deviation) or frequency. Abbreviations: ADNI, Alzheimer’s Disease Neuroimaging Initiative; BLSA, Baltimore Longitudinal Study of Aging; VMAP, Vanderbilt Memory & Aging Project; yrs, years; *APOE*-ε4, apolipoprotein ε4; *APOE*-ε2, apolipoprotein ε2. \*In participants at least 2 visits.

### dMRI Acquisition and Preprocessing

All data were preprocessed using the *PreQual* pipeline, which is an automated pipeline that corrects diffusion MRI data for distortions/motion and eddy currents^49, 50^. The quality control PDFs which are outputted by the *PreQual* pipeline were manually inspected, and participants with poor quality data were excluded from analysis. This data was then inputted into DTIFIT to calculate conventional (i.e., uncorrected) diffusion MRI metrics, including fractional anisotropy (FA_CONV_), mean diffusivity (MD_CONV_), axial diffusivity (AxD_CONV_), and radial diffusivity (RD_CONV_). The preprocessed data were also inputted into MATLAB code to calculate free-water (FW) corrected metrics^36^, including FW-corrected fractional anisotropy (FA_FWcorr_), FW-corrected mean diffusivity (MD_FWcorr_), FW-corrected axial diffusivity (AxD_FWcorr_), and FW-corrected radial diffusivity (RD_FWcorr_)^36^. A standard space representation of these maps was created by non-linearly registering the FA_CONV_ map to the FMRIB58_FA atlas using the Advanced Normalization Tools (ANTs) package^51^. The warp obtained from this registration was then applied to all other microstructural maps.

### White Matter Tractography Templates

All tractography templates used in this study were drawn from existing resources^17, 22, 32–35^ and are available in a publicly available GitHub repository (https://github.com/VUMC-VMAC/Tractography_Templates). In total, this study used 48 white matter tractography templates (**Figure 1A**) which can be grouped into 7 different tract-types, including association (n=3), limbic (n=7), projection (n=9), motor transcallosal (TC) (n=6), occipital TC (n=6), parietal TC (n=5), and prefrontal TC (n=12) tracts. The association tracts included the inferior fronto-occipital fasciculus (IFOF), superior longitudinal fasciculus (SLF), and the SLF temporoparietal (SLF-TP) tracts. The limbic tracts included the inferior longitudinal fasciculus (ILF), cingulum bundle, fornix, uncinate fasciculus (UF), superior temporal gyrus TC, middle temporal gyrus TC, and inferior temporal gyrus tracts. The projection tracts included several descending sensorimotor tracts, including projections from the somatosensory cortex (S1), primary motor cortex (M1), supplemental motor area (SMA), dorsal premotor cortex (PMd), ventral premotor cortex (PMv), and pre-supplemental motor area (preSMA). Additionally, projections tracts from the superior frontal gyrus to caudate, middle frontal gyrus to caudate, and M1 to putamen (corticostriatal) were included. The motor TC tracts included homologous connections of the S1, M1, PMd, PMv, SMA, and preSMA tracts. The occipital TC tracts included homologous connections of the lingual gyrus, superior occipital gyrus, middle occipital gyrus, inferior occipital gyrus, cuneus, and calcarine sulcus. The parietal TC tracts included homologous connections of the superior parietal lobe (SPL), inferior parietal lobe (IPL), paracentral lobe, angular gyrus, and supramarginal gyrus. The prefrontal TC tracts included homologous connections of the superior frontal gyrus, medial frontal gyrus, middle frontal gyrus, inferior frontal gyrus (IFG) pars opercularis, IFG pars triangularis, IFG pars orbitalis, lateral orbital gyrus, medial orbital gyrus, anterior orbital gyrus, medial orbitofrontal gyrus, gyrus rectus, and olfactory cortex.

**Figure 1.**
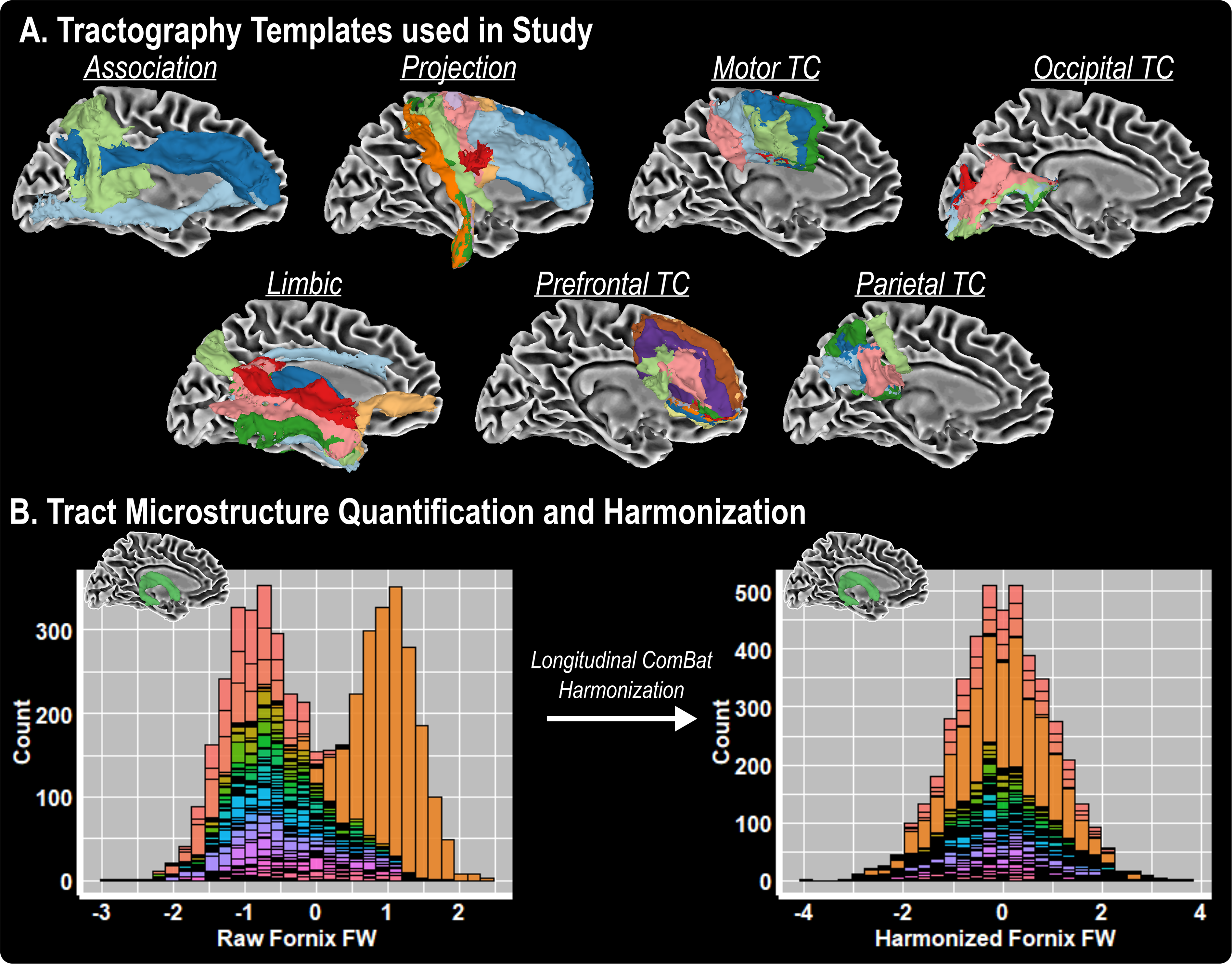
Tractography templates used in study. 48 white matter tractography templates (**A**) were used to evaluate aging and these tracts can be grouped into 7 different tract-types, including association, projection, motor transcallosal (TC), occipital TC, limbic, prefrontal TC, and parietal TC tracts. *Longitudinal ComBat* harmonization was conducted on all imaging features to harmonize across all *site x scanner x protocol* combinations – raw and harmonized fornix FW values are shown and colored by each *site x scanner x protocol* combination (**B**).

### Diffusion MRI Data Harmonization

A region of interest approach was used to calculate mean conventional (FA_CONV_, MD_CONV_, AxD_CONV_, RD_CONV_) and FW-corrected (FW, FA_FWcorr_, MD_FWcorr_, AxD_FWcorr_, RD_FWcorr_) microstructure within all tractography templates for each participant, resulting in 432 unique values for each imaging session. These values were subsequently harmonized using the *Longitudinal ComBat* technique in R (version 4.1.0)^52^. In the *Longitudinal ComBat* harmonization, a batch variable which controlled for all *site x scanner x protocol* combinations was used. We also used several covariates to control for between-scanner, between-protocol, and between-cohort effects, including mean-centered age, mean-centered age squared, education, race/ethnicity, diagnosis at baseline, *APOE*-ε4 positivity, *APOE*-ε2 positivity, the interaction of age and aging type (i.e., normal, abnormal), and the interaction of mean-centered age and aging type. Leveraging the *Longitudinal ComBat* harmonization technique, we were also able to model the random effects of aging for each participant [i.e., ∼1+age|participant]. **Figure 1B** illustrates the raw and harmonized values for fornix FW. The harmonized values were then scaled by their standard deviation and used in all subsequent statistical analyses.

### Statistical Analyses

All statistical analyses were performed in R (version 4.1.0), and age was mean centered prior to analysis. First, stratified (normal aging or abnormal aging) linear mixed effects (LME) regression analysis was conducted on all 9 diffusion MRI metrics to determine the effect of aging within the white matter tractography templates. Fixed effects in stratified models included age, age squared, education, sex, race/ethnicity, *APOE*-ε4 positivity, and *APOE*-ε2 positivity. Random effects included intercept and age. Separate LMEs were created for all 9 metrics across all 48 white matter tracts, resulting in 432 models. For stratified models, the effect of aging was evaluated by focusing on the statistics from the *age* term. Following stratified analysis, similar LMEs were built for the entire cohort, adding *age x cognitive status* and *age*^2^ *x cognitive status* interaction terms as well as all lower order terms. For the interaction analysis, we focused on the statistics from the *age x cognitive status* term. Significance was set a prior as α=0.05 and correction for multiple comparisons was made using the false discovery rate method.

Follow-up bootstrapped (n=1000 for each microstructural measure x tract combination) LME analyses were also conducted to determine which microstructural measures were most vulnerable to abnormal aging. For each microstructural variable, a base LME model covaried for age, age^2^, education, sex, race/ethnicity, *APOE*-ε4 positivity, and *APOE*-ε2 positivity, whereas a more comprehensive model included the previous covariates plus *age x cognitive status* and *age*^2^ *x cognitive status* interaction terms and all lower order terms. Marginal variance (i.e., the variance of the fixed effects) was pulled from all respective models and differences between the comprehensive model and base model were quantified. A repeated measures ANOVA was then conducted to compare the effect of microstructure on difference in marginal variance, using each of the tracts as a pairwise variable. Follow-up one way ANOVAs were then conducted for each microstructural variable to determine which tracts had the most significant differences in marginal variance between models.

## Results

### White Matter Decline in Normal and Abnormal Aging

The effects of normal aging on conventional and FW-corrected metrics are shown in the heatmap in **Figure 2** and relevant statistics for all normal aging effects can be found in **Supplemental Table 2**. As shown in **Figure 2**, there was a near global effect of normal aging on white matter microstructure. Tracts which were most sensitive to aging included the fornix (**Figure 2A**), TC inferior frontal gyrus (IFG) pars opercularis (**Figure 2B**), caudate to middle frontal (**Figure 2C**), and TC middle frontal gyrus (**Figure 2D**) tracts. The effects of abnormal aging on conventional and FW-corrected metrics are shown in the heatmap in **Figure 3** and relevant statistics for all abnormal aging effects can be found in **Supplemental Table 3**. As shown in **Figure 3**, there was a near global effect of abnormal aging on white matter microstructure, and the top involved tracts included the inferior frontal occipital fasciculus (IFOF) (**Figure 3A**), caudate to middle frontal (**Figure 3B**), fornix (**Figure 3C**), and TC IFG pars opercularis (**Figure 3D**) tracts.

**Figure 2.**
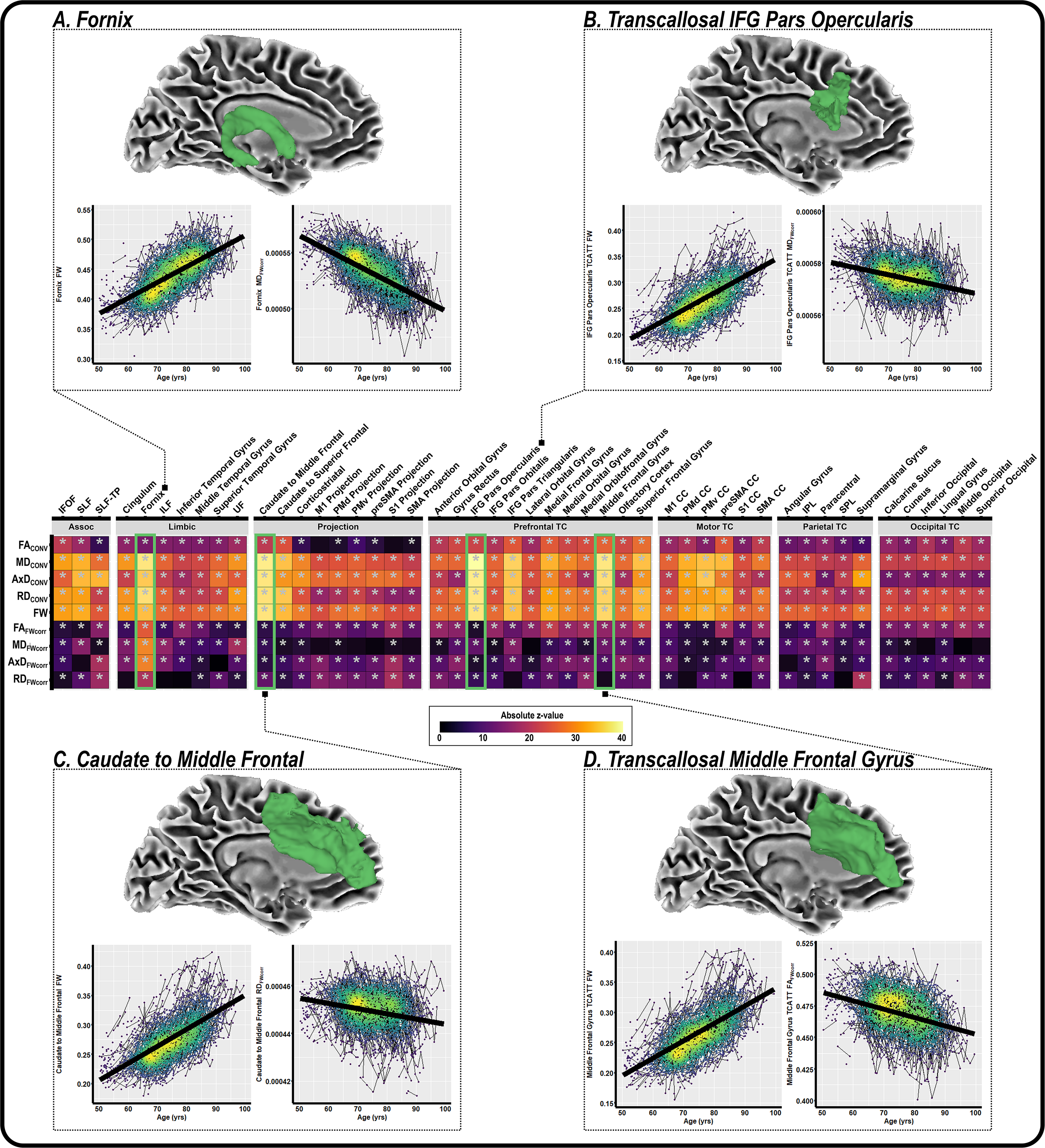
The effect of normal aging on white matter microstructure. Linear mixed effects (LME) regression was conducted for each conventional (MD_CONV_, FA_CONV_, AxD_CONV_, RD_CONV_) and FW-corrected measure (FW, MD_FWcorr_, FA_FWcorr_, AxD_FWcorr_, RD_FWcorr_) to determine the association of normal aging with white matter microstructure. The heatmap, grouped by tract-type, illustrates the t-value for each independent LME regression. Blocks marked with an asterisk represent associations meeting the p_FDR_<0.05 threshold. Examples for the normal aging effect on white matter microstructure are shown for the fornix (**A**), transcallosal IFG pars opercularis (**B**), caudate to middle frontal gyrus (**C**), and transcallosal middle frontal gyrus (**D**) tracts.

**Figure 3.**
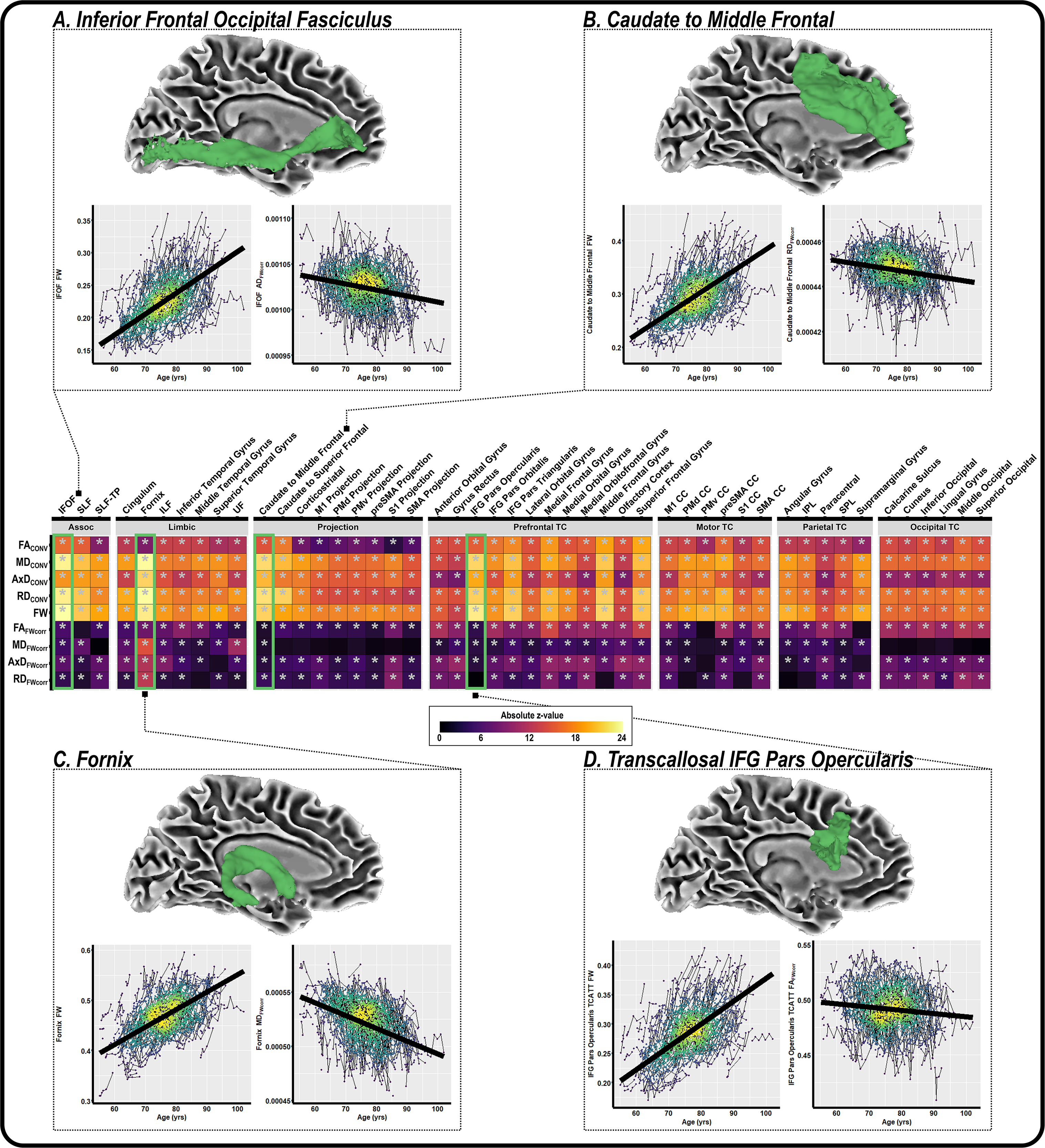
The effect of abnormal aging on white matter microstructure. Linear mixed effects (LME) regression was conducted for each conventional (MD_CONV_, FA_CONV_, AxD_CONV_, RD_CONV_) and FW-corrected measure (FW, MD_FWcorr_, FA_FWcorr_, AxD_FWcorr_, RD_FWcorr_) to determine the association of abnormal aging with white matter microstructure. The heatmap, grouped by tract-type, illustrates the t-value for each independent LME regression. Blocks marked with an asterisk represent associations meeting the p_FDR_<0.05 threshold. Examples for the abnormal aging effect on white matter microstructure are shown for the inferior frontal occipital fasciculus (**A**), caudate to middle frontal (**B**), fornix (**C**), and transcallosal inferior frontal gyrus (IFG) pars opercularis (**D**) tracts.

### Differential White Matter Decline in Normal and Abnormal Aging

The interactions between normal and abnormal aging on conventional and FW-corrected metrics are shown in the heatmap in **Figure 4** and all relevant statistics can be found in **Supplemental Table 4**. Several of the top interactions are illustrated in **Figures 4A-D**, including IFOF FW (β=-3.63x10^-3^, p_FDR_=1.36x10^-6^, z=-5.59) in **Figure 4A**, TC IFG pars orbitalis FW (β=-4.12x10^-3^, p_FDR_=1.53x10^-7^, z=-6.07) in **Figure 4B**, fornix FA_FWcorr_ (β=-1.36x10^-3^, p_FDR_=3.93x10^-8^, z=-6.39) in **Figure 4C**, and TC angular gyrus FW (β=-4.30x10^-3^, p_FDR_=6.02x10^-7^, z=-5.78) in **Figure 4D**.

**Figure 4.**
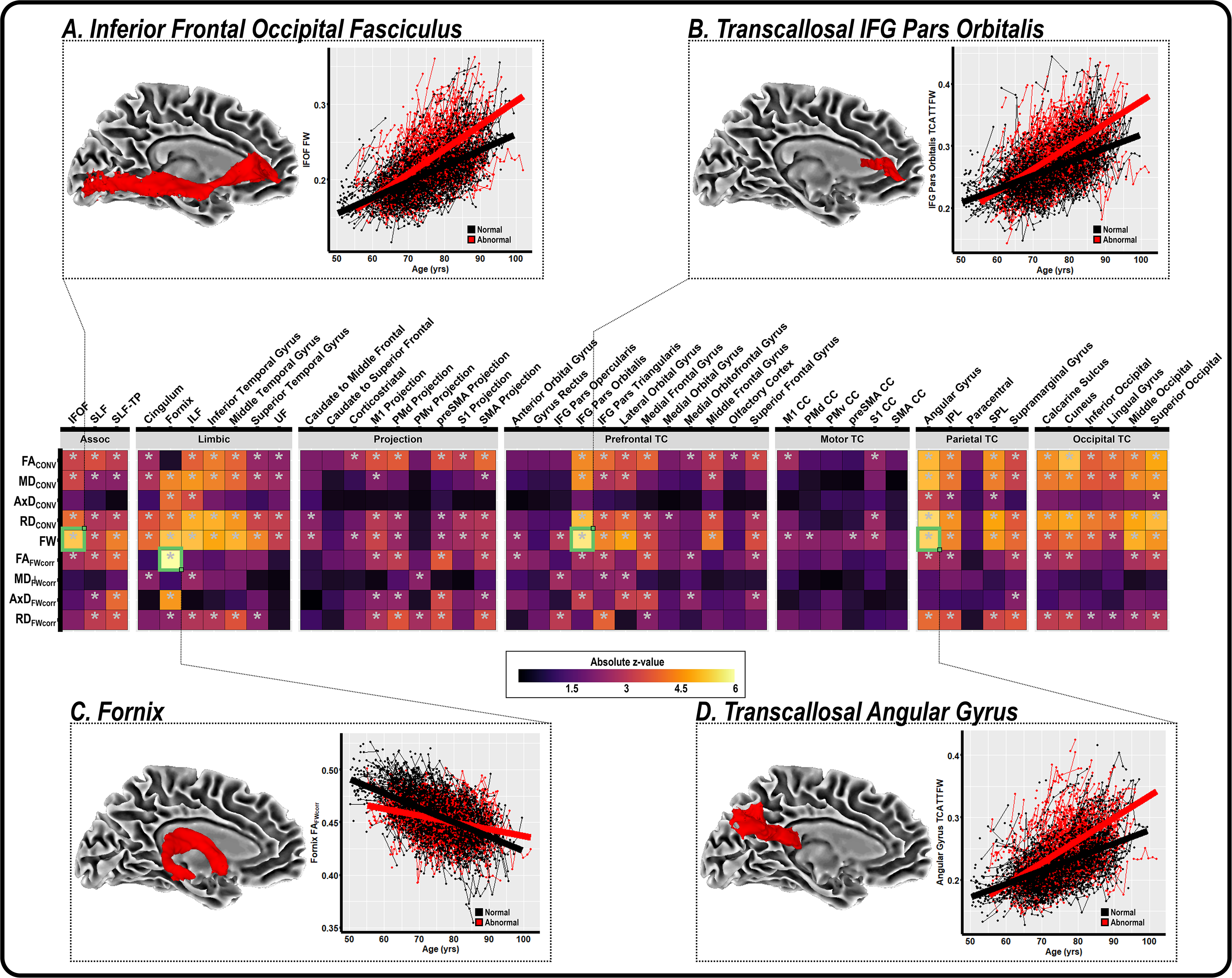
Differential changes in white matter microstructure in normal aging versus cognitively impaired groups. Linear mixed effects (LME) regression was conducted for each conventional (MD_CONV_, FA_CONV_, AxD_CONV_, RD_CONV_) and FW-corrected measure (FW, MD_FWcorr_, FA_FWcorr_, AxD_FWcorr_, RD_FWcorr_) to quantify the *age x cognitive status* interaction effect. The heatmap, grouped by tract-type, illustrates the t-value for each independent LME regression. Blocks marked with an asterisk represent associations meeting the p_FDR_<0.05 threshold. Examples for the *age x cognitive status* interaction effect on white matter microstructure are shown for the inferior frontal occipital fasciculus (**A**), transcallosal IFG pars orbitalis (**B**), fornix (**C**), and transcallosal angular gyrus (**D**) tracts.

### Bootstrapped Analysis to Determine White Matter Microstructural Measures Most Vulnerable to Abnormal Aging

A bootstrapped analysis was conducted to determine the microstructural measures and tract most vulnerable to abnormal aging. A repeated measures ANOVA, controlling for tract, was significant (p<0.05), and post-hoc analyses found that the FW measure was most sensitive to abnormal aging. **Figure 5A** illustrates the mean ΔR^2^ for FW within all tracts. **Figure 5B** shows the mean and standard error of the bootstrapped ΔR^2^ for all tracts for the FW measure. An ANOVA was conducted to determine if there were significant differences in ΔR^2^ for each tract and results were significant (p<0.05). Post-hoc analyses were conducted to determine which tracts were most vulnerable to abnormal aging, and we found that the cingulum bundle was most vulnerable to abnormal aging. Illustrations of the top 6 vulnerable tracts to abnormal aging are shown in **Figure 5C**. All pairwise comparisons for FW can be found in **Supplemental Table 5**.

**Figure 5.**
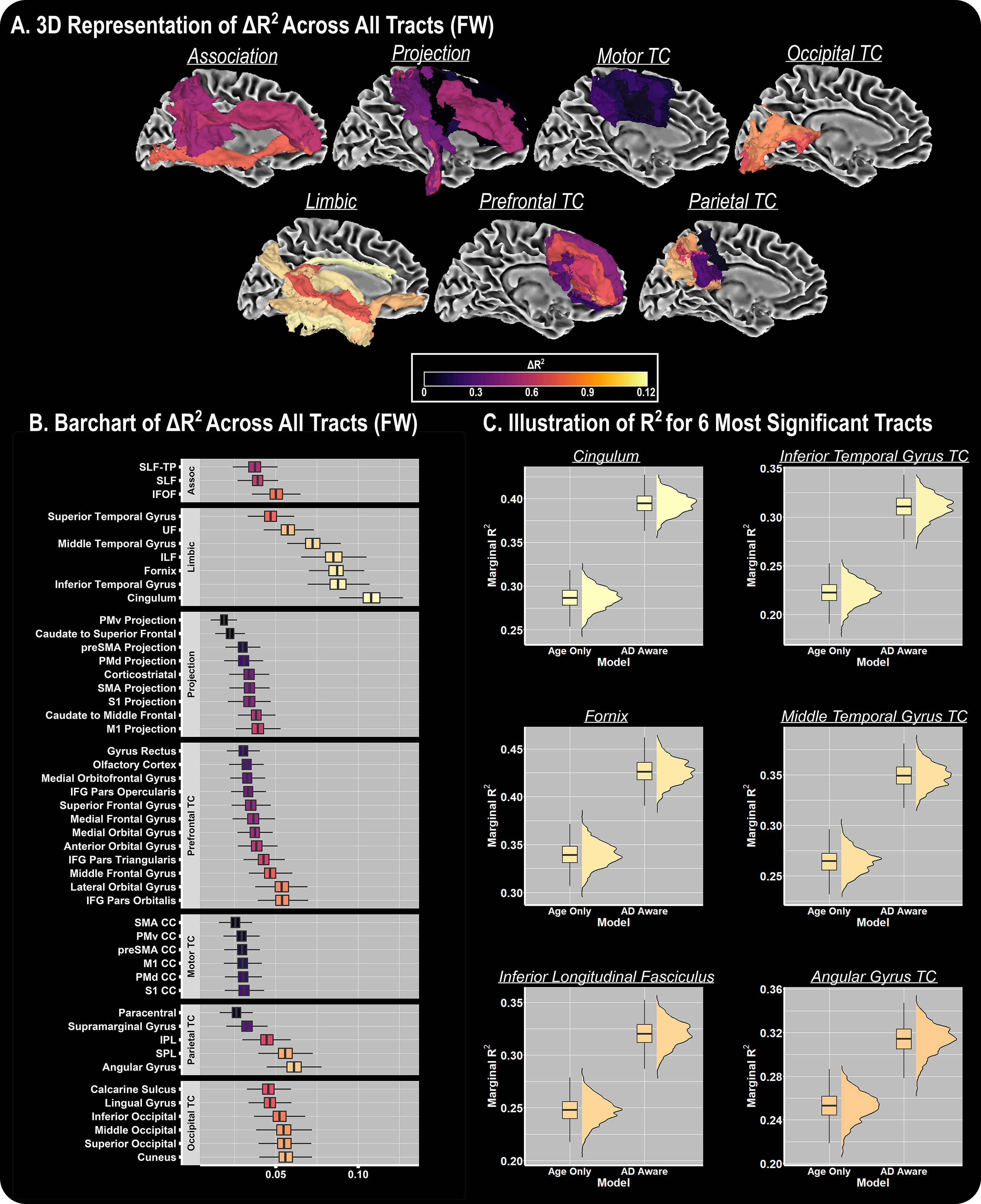
FW Vulnerability to Abnormal Aging. Mean ΔR^2^ between the base model (i.e., only age and covariates) and AD aware (i.e., includes abnormal aging covariates) models for FW is shown on all white matter tract templates (**A**), and a bar chart summarizes the mean and standard deviation for all bootstrapped differences between models (**B**). Base and comprehensive model marginal R^2^ values are shown for the 6 most significant tracts.

## Discussion

We used a longitudinal, multi-site set of cohorts to examine the relationship of normal and abnormal aging with white matter microstructural decline. Specifically, we conducted robust preprocessing in 1,723 participants across 4,605 imaging sessions to create conventional and FW corrected dMRI maps. We then quantified 4 conventional (FA_CONV_, AxD_CONV_, RD_CONV_, MD_CONV_) and 5 FW corrected (FA_FWcorr_, AxD_FWcorr_, RD_FWcorr_, MD_FWcorr_, FW) microstructural values within 48 well-established white matter tractography templates spanning the association, limbic, projection, motor transcallosal (TC), occipital TC, parietal TC, and prefrontal TC tracts. Subsequently, all microstructural values (n=432) were inputted into the *Longitudinal ComBat* toolkit to harmonize between all *site x scanner x protocol* combinations (n=108). The harmonized microstructural values were used to: (1) evaluate the effects of normal aging on white matter microstructural decline, (2) evaluate the effects of abnormal aging on white matter microstructural decline, (3) quantify differential magnitudes of white matter microstructural decline in the normal aging versus the abnormal aging group, and (4) conduct robust bootstrapping analysis to determine which tracts are most vulnerable in abnormal aging. Through these targeted analyses, we replicated prior studies by demonstrating that there is a global association between age and white matter microstructural decline, with pronounced effects within the limbic and prefrontal transcallosal tracts. Further, we found several interactions, indicating steeper rates of white matter decline in the abnormal compared with abnormal group, with the most significant interactions being within the fornix, transcallosal IFG pars orbitalis, angular gyrus, and inferior frontal occipital fasciculus. Finally, bootstrapped analyses found that the FW measure is most vulnerable to abnormal aging, with the limbic tracts (e.g., cingulum bundle) demonstrating enhanced vulnerability. Together, these results suggest that the FW measure is highly sensitive in both normal and abnormal aging and is particularly sensitive to detecting microstructural decline in neurodegenerative disease.

### White Matter Microstructural Decline in Normal Aging

The present study provides novel findings to a long-standing line of research focusing on the relationship between normal aging (n=1,155) and white matter microstructural decline^1–15^. Using both conventional and FW corrected dMRI metrics in conjunction with spatially precise white matter tractography templates, we found global associations with normal aging. We found that the most pronounced effects were within prefrontal transcallosal, prefrontal projection, and limbic tracts. For instance, we found that that normal aging was associated with pronounced microstructural abnormalities in the fornix, transcallosal IFG pars opercularis, caudate to middle frontal gyrus, and transcallosal middle frontal gyrus tracts. These results corroborate prior studies which suggest that the prefrontal tracts projecting from the genu of the corpus callosum are sensitive to aging^7, 8, 10, 20^. Moreover, this adds further support to the “last-in-first-out” paradigm, which suggests that tracts which take the longest to develop are the ones which are most vulnerable to age-related processes^15^. Prior studies have also shown that the association tracts, such as the inferior frontal occipital fasciculus (IFOF), are involved in the aging process. For example, *Cox et al.* demonstrated that the older age was associated with lower white matter microstructural values in the association tracts^53^. Importantly, we replicate these findings and demonstrate that IFOF FW is the most significantly associated microstructural metric with normal aging. Nevertheless, the effect sizes in many of the prefrontal transcallosal, prefrontal projection, and limbic tracts were stronger than the association tracts.

### White Matter Microstructural Decline in Abnormal Aging

While several prior publications have evaluated normal aging, far fewer studies have leveraged large-scale dMRI data to quantify the effects of abnormal aging and white matter microstructural decline. In our study of participants with cognitive impairment (n=568), we found global associations between aging and white matter microstructural decline, with the association, limbic, prefrontal projection, and prefrontal transcallosal tracts having the most sensitive associations. The most sensitive association in the analysis of the cognitively impaired subgroup was for IFOF FW, in which aging was associated with higher IFOF FW. While longitudinal studies of white matter in cognitively impaired samples are sparse, our results are comparable to cross-sectional studies of white matter microstructure along the AD continuum, which suggest that posterior, temporal, and prefrontal tracts are most involved in AD white matter microstructural decline^4, 14, 23–29^.

### Differences between Normal and Abnormal Aging

Although main effects of normal and abnormal aging on white matter microstructural decline are fundamental to our understanding of the aging process, it is essential to understand how aging differs between participants who do and do not progress along the spectrum of cognitive impairment. Our interaction analyses found that several white matter microstructural metrics interact with normal and abnormal aging groups, with strong effects in the IFOF, transcallosal IFG pars orbitalis, fornix, and transcallosal angular gyrus. The strongest interaction was found for fornix FA_FWcorr_, in which abnormal agers had a lower overall FA_FWcorr_ earlier in age. In this analysis, we found that FW was especially sensitive to detecting interactions between normal and abnormal agers. Strong FW interactions were found for the IFOF, transcallosal IFG pars orbitalis, and transcallosal angular gyrus. These findings are consistent with a prior study demonstrating that IFOF has more rapid decline (i.e., decrease in FA_CONV_) in individuals with subsequent cognitive impairment compared to normal agers^31^. We corroborate these findings while simultaneously demonstrating that the FW metric may be more sensitive than FA_CONV_. Further, this is the first study to demonstrate – in a comprehensive multi-site cohort – that abnormal agers have differential patterns of white matter microstructural decline.

To further understand the vulnerability of white matter microstructure to abnormal aging, bootstrapping analyses were conducted, in which we compared the marginal R^2^ in a model only covarying for age with a model covarying for age and cognitive status (normal/abnormal). From this analysis, we found that the FW measure is the most vulnerable to abnormal aging. Post-hoc pairwise t-tests were conducted to determine which tracts were most vulnerable to abnormal aging for the FW metric, in which we found that limbic tracts were most vulnerable. When comparing limbic tracts, we found that the cingulum bundle was the most vulnerable, followed closely by the transcallosal inferior temporal gyrus, fornix, and inferior longitudinal fasciculus. Together, our results suggest that FW -- and more specifically FW within the limbic tracts – should be further studied to enhance our knowledge into the differences in normal and abnormal aging.

### Strengths/Weaknesses

The current study has several strengths, including a harmonized multi-site diffusion MRI cohort which far exceeds the sample size in any previous single-shell FW study on aging (n=1,723). An additional strength of this study is that we used longitudinal data with a total of 4,605 sessions. While several prior studies have evaluated white matter neurodegeneration in aging, many of them have used long-standing tractography templates (e.g., JHU template) or regions of interest within specific white matter areas (e.g., genu and splenium of the corpus callosum). A major novelty of this study is that we have used 48 recently developed tractography templates spanning the association, limbic, projection, and transcallosal tracts. For this reason, we were able to capture specificity in white matter neurodegenerative aging in the prefrontal cortex which was not previously known. Importantly, these tractography templates are freely accessible (https://github.com/VUMC-VMAC/Tractography_Templates). We hope that providing these templates will encourage replication studies and further studies evaluating neurodegeneration in aging. Finally, a major strength of this study is that it used the single-shell FW technique – this technique (and others) allows researchers to apply advanced post-processing techniques to long-standing cohorts of aging (e.g., ADNI, BLSA, VMAP) which may not always have multi-shell diffusion MRI acquisition for all participants. While our bootstrapped analyses suggest that FW is the most sensitive measure to abnormal aging, conventional dMRI measures also showed high sensitivity to abnormal aging. Therefore, the use of this measure may not be necessary if the goal is classification of abnormal aging; however, if the goal is to understand which biological mechanisms are associated with abnormal aging, FW correction may provide additional insight. Despite these strengths, this study comprised participants who were predominantly well-educated, non-Hispanic white individuals. For this reason, our results may not be generalizable to other cohorts. An additional limitation of our study is that diffusion MRI was obtained from several sites, scanners, and with several different scanning protocols. While we used harmonization protocols to account for this (i.e., *Longitudinal ComBat*), there is no technique which will completely mitigate the heterogeneity provided by these variables. Finally, we did not pair our neuroimaging analyses with any biomarkers, therefore limiting our ability to tie age-related neurodegeneration to specific biologic pathways. Future studies which use comparable sample sizes and incorporate biomarkers of white matter neurodegeneration (e.g., neurofilament light) and astrocytosis may further enhance our understanding of the biology underpinning the white matter microstructural changes seen in aging.

In conclusion, this multi-site longitudinal study provides strong evidence that normal and abnormal aging are both associated with white matter microstructural decline, and that the limbic tracts are most affected in abnormal aging. Therefore, we suggest that FW correction be conducted when using single-shell diffusion MRI acquisition scans to evaluate aging. Future large-scale analyses should pair dMRI metrics with biomarkers of disease to help further understand the biological processes associated with white matter microstructural decline in abnormal aging.

## Appendix 1. Collaborators

*Data used in preparation of this article were obtained from the Alzheimer’s Disease Neuroimaging Initiative (ADNI) database (adni.loni.usc.edu). As such, the investigators within the ADNI contributed to the design and implementation of ADNI and/or provided data but did not participate in analysis or writing of this report. A complete listing of ADNI investigators can be found at: https://adni.loni.usc.edu/wp-content/uploads/how_to_apply/ADNI_Acknowledgement_List.pdf

## Literature Cited

1. Bennett, I. J., Madden, D. J., Vaidya, C. J., Howard, D. V. & Howard Jr., J. H. Age-related differences in multiple measures of white matter integrity: A diffusion tensor imaging study of healthy aging. Human Brain Mapping 31, 378–390 (2010).

2. Bender, A. R. & Raz, N. Normal-appearing cerebral white matter in healthy adults: mean change over 2 years and individual differences in change. Neurobiology of Aging 36, 1834–1848 (2015).

3. Brickman, A. M. et al. Testing the white matter retrogenesis hypothesis of cognitive aging. Neurobiology of Aging 33, 1699–1715 (2012).

4. Sexton, C. E., Kalu, U. G., Filippini, N., Mackay, C. E. & Ebmeier, K. P. A meta-analysis of diffusion tensor imaging in mild cognitive impairment and Alzheimer’s disease. Neurobiol Aging 32, 2322 e5–18 (2011).

5. Lebel, C. et al. Diffusion tensor imaging of white matter tract evolution over the lifespan. Neuroimage 60, 340–52 (2012).

6. Gazes, Y. et al. White matter tract covariance patterns predict age-declining cognitive abilities. NeuroImage 125, 53–60 (2016).

7. Salat, D. H. et al. Age-related alterations in white matter microstructure measured by diffusion tensor imaging. Neurobiology of Aging 26, 1215–1227 (2005).

8. Davis, S. W. et al. Assessing the effects of age on long white matter tracts using diffusion tensor tractography. NeuroImage 46, 530–541 (2009).

9. Lövdén, M. et al. The dimensionality of between-person differences in white matter microstructure in old age. Human Brain Mapping 34, 1386–1398 (2013).

10. Kochunov, P. et al. Relationship between white matter fractional anisotropy and other indices of cerebral health in normal aging: tract-based spatial statistics study of aging. Neuroimage 35, 478–87 (2007).

11. Westlye, L. T. et al. Life-span changes of the human brain white matter: diffusion tensor imaging (DTI) and volumetry. Cerebral cortex 20, 2055–2068 (2010).

12. Yeatman, J. D., Wandell, B. A. & Mezer, A. A. Lifespan maturation and degeneration of human brain white matter. Nat Commun 5, 4932 (2014).

13. de Groot, M. et al. White Matter Degeneration with Aging: Longitudinal Diffusion MR Imaging Analysis. Radiology 279, 532–541 (2016).

14. Stricker, N. H. et al. Decreased white matter integrity in late-myelinating fiber pathways in Alzheimer’s disease supports retrogenesis. NeuroImage 45, 10–16 (2009).

15. Kiely, M. et al. Insights into human cerebral white matter maturation and degeneration across the adult lifespan. NeuroImage 247, 118727 (2022).

16. 2021 Alzheimer’s disease facts and figures. Alzheimers Dement 17, 327–406 (2021).

17. Archer, D. B. et al. Free-water metrics in medial temporal lobe white matter tract projections relate to longitudinal cognitive decline. Neurobiol Aging 94, 15–23 (2020).

18. Archer, D. B. et al. The relationship between white matter microstructure and self-perceived cognitive decline. NeuroImage: Clinical 32, 102794 (2021).

19. Blanchard, J. W. et al. APOE4 impairs myelination via cholesterol dysregulation in oligodendrocytes. Nature 611, 769–779 (2022).

20. Bartzokis, G. et al. Heterogeneous age-related breakdown of white matter structural integrity: implications for cortical “disconnection” in aging and Alzheimer’s disease. Neurobiology of Aging 25, 843–851 (2004).

21. Mori, S., Wakana, S., Zijl, P. C. M. van & Nagae-Poetscher, L. M. MRI Atlas of Human White Matter. (Elsevier, 2005).

22. Mori, S. et al. Stereotaxic white matter atlas based on diffusion tensor imaging in an ICBM template. NeuroImage 40, 570–582 (2008).

23. Walsh, R., Bergamino, M. & Stokes, A. Free-Water Diffusion Tensor Imaging (DTI) Improves the Accuracy and Sensitivity of White Matter Analysis in Alzheimer’s Disease (4979). Neurology 94, (2020).

24. Berlot, R., Metzler-Baddeley, C., Jones, D. K. & O’Sullivan, M. J. CSF contamination contributes to apparent microstructural alterations in mild cognitive impairment. NeuroImage 92, 27–35 (2014).

25. Dumont, M. et al. Free Water in White Matter Differentiates MCI and AD From Control Subjects. Frontiers in Aging Neuroscience 11, (2019).

26. Schouten, T. M. et al. Individual classification of Alzheimer’s disease with diffusion magnetic resonance imaging. NeuroImage 152, 476–481 (2017).

27. Dalboni da Rocha, J. L., Bramati, I., Coutinho, G., Tovar Moll, F. & Sitaram, R. Fractional Anisotropy changes in Parahippocampal Cingulum due to Alzheimer’s Disease. Sci Rep 10, 2660 (2020).

28. Bozzali, M. et al. Damage to the cingulum contributes to Alzheimer’s disease pathophysiology by deafferentation mechanism. Hum Brain Mapp 33, 1295–308 (2012).

29. Nir, T. M. et al. Effectiveness of regional DTI measures in distinguishing Alzheimer’s disease, MCI, and normal aging. NeuroImage: Clinical 3, 180–195 (2013).

30. Nemy, M. et al. Cholinergic white matter pathways along the Alzheimer’s disease continuum. Brain awac385 (2022) doi:10.1093/brain/awac385.

31. Shafer, A. T. et al. Accelerated decline in white matter microstructure in subsequently impaired older adults and its relationship with cognitive decline. Brain Communications 4, fcac051 (2022).

32. Brown, C. A. et al. Development, validation and application of a new fornix template for studies of aging and preclinical Alzheimer’s disease. Neuroimage Clin 13, 106–115 (2017).

33. Archer, D. B., Vaillancourt, D. E. & Coombes, S. A. A Template and Probabilistic Atlas of the Human Sensorimotor Tracts using Diffusion MRI. Cereb Cortex 28, 1685–1699 (2018).

34. Archer, D. B. et al. Development and validation of the automated imaging differentiation in parkinsonism (AID-P): a multicentre machine learning study. The Lancet Digital Health 1, e222–e231 (2019).

35. Archer, D. B., Coombes, S. A., McFarland, N. R., DeKosky, S. T. & Vaillancourt, D. E. Development of a transcallosal tractography template and its application to dementia. Neuroimage 200, 302–312 (2019).

36. Pasternak, O., Sochen, N., Gur, Y., Intrator, N. & Assaf, Y. Free water elimination and mapping from diffusion MRI. Magn Reson Med 62, 717–30 (2009).

37. Zhang, H., Schneider, T., Wheeler-Kingshott, C. A. & Alexander, D. C. NODDI: practical in vivo neurite orientation dispersion and density imaging of the human brain. Neuroimage 61, 1000–16 (2012).

38. Archer, D. B., Patten, C. & Coombes, S. A. Free-water and free-water corrected fractional anisotropy in primary and premotor corticospinal tracts in chronic stroke. Hum Brain Mapp 38, 4546– 4562 (2017).

39. Archer, D. B. et al. A widespread visually-sensitive functional network relates to symptoms in essential tremor. Brain 141, 472–485 (2018).

40. Burciu, R. G. et al. Progression marker of Parkinson’s disease: a 4-year multi-site imaging study. Brain 140, 2183–2192 (2017).

41. Yang, J. et al. Multimodal dopaminergic and free-water imaging in Parkinson’s disease. Parkinsonism Relat Disord 62, 10–15 (2019).

42. Ofori, E. et al. Longitudinal changes in free-water within the substantia nigra of Parkinson’s disease. Brain 138, 2322–2331 (2015).

43. Carreira Figueiredo, I., Borgan, F., Pasternak, O., Turkheimer, F. E. & Howes, O. D. White-matter free-water diffusion MRI in schizophrenia: a systematic review and meta-analysis. Neuropsychopharmacol. 47, 1413–1420 (2022).

44. Ofori, E. et al. Free-water imaging of the hippocampus is a sensitive marker of Alzheimer’s disease. Neuroimage Clin 24, 101985 (2019).

45. Chad, J. A., Pasternak, O., Salat, D. H. & Chen, J. J. Re-examining age-related differences in white matter microstructure with free-water corrected diffusion tensor imaging. Neurobiology of Aging 71, 161–170 (2018).

46. Ferrucci, L. The Baltimore Longitudinal Study of Aging (BLSA): A 50-Year-Long Journey and Plans for the Future. The Journals of Gerontology: Series A 63, 1416–1419 (2008).

47. Jack, C. R. et al. The Alzheimer’s disease neuroimaging initiative (ADNI): MRI methods. Journal of Magnetic Resonance Imaging 27, 685–691 (2008).

48. Jefferson, A. L. et al. The Vanderbilt Memory & Aging Project: Study Design and Baseline Cohort Overview. Journal of Alzheimer’s Disease 1–20 (2016).

49. Cai, L. Y. et al. PreQual: An automated pipeline for integrated preprocessing and quality assurance of diffusion weighted MRI images. Magnetic Resonance in Medicine 86, 456–470 (2021).

50. Schilling, K. G. et al. Synthesized b0 for diffusion distortion correction (Synb0-DisCo). Magn Reson Imaging 64, 62–70 (2019).

51. Avants, B. B., Epstein, C. L., Grossman, M. & Gee, J. C. Symmetric diffeomorphic image registration with cross-correlation: evaluating automated labeling of elderly and neurodegenerative brain. Med Image Anal 12, 26–41 (2008).

52. Beer, J. C. et al. Longitudinal ComBat: A method for harmonizing longitudinal multi-scanner imaging data. NeuroImage 220, 117129 (2020).

53. Cox, S. R. et al. Ageing and brain white matter structure in 3,513 UK Biobank participants. Nat Commun 7, 13629 (2016).

